# Effects of cell cycle variability on lineage and population measurements of mRNA abundance

**DOI:** 10.1101/2020.03.24.006494

**Authors:** Ruben Perez-Carrasco, Casper Beentjes, Ramon Grima

## Abstract

Many models of gene expression do not explicitly incorporate a cell cycle description. Here we derive a theory describing how mRNA fluctuations for constitutive and bursty gene expression are influenced by stochasticity in the duration of the cell cycle and the timing of DNA replication. Analytical expressions for the moments show that omitting cell cycle duration introduces an error in the predicted mean number of mRNAs that is a monotonically decreasing function of *η*, which is proportional to the ratio of the mean cell cycle duration and the mRNA lifetime. By contrast, the error in the variance of the mRNA distribution is highest for intermediate values of *η* consistent with genome-wide measurements in many organisms. Using eukaryotic cell data, we estimate the errors in the mean and variance to be at most 3% and 25%, respectively. Furthermore, we derive an accurate negative binomial mixture approximation to the mRNA distribution. This indicates that stochasticity in the cell cycle can introduce fluctuations in mRNA numbers that are similar to the effect of bursty transcription. Finally, we show that for real experimental data, disregarding cell cycle stochasticity can introduce errors in the inference of transcription rates larger than 10%.

## Introduction

Intrinsic noise in gene expression induces variability in the transcript number across a population of cells. Current microscopy techniques are able to capture this variability, which can be used to infer the kinetic parameters of transcription, thereby letting us quantify mechanisms in charge of the regulation of gene expression [1–3]. In order to make this inference possible, it is necessary to have an accurate stochastic dynamical model that is able to relate the details of the mRNA number distribution to the different transcriptional and post-transcriptional molecular mechanisms involved in mRNA processing. This has been extensively done by describing the dynamics of the system by means of the Master Equation, a Markovian description whose solution gives the probability of observing a certain number of mRNAs in a cell at a certain time [4]. Since the exact analytical solution of the Master Equation is only available for a few scenarios (e.g. [5–7]), the study of the probability distribution of mRNA transcript number is usually limited to calculating the moments of the distribution.

One particular mechanism that has been difficult to study analytically is the influence of the cell cycle on the distribution of mRNAs in a population of cells. The duration of the different phases of the cell cycle is stochastic, introducing noise not only in the time of mitosis when the molecular content is diluted but also in the time at which DNA is replicated, which in turn increases the mRNA production rate [3]. In addition, during mitosis, the cellular content is divided, leading to a stochastic transcript bipartition [8].

Due to these different challenges, mathematical effort has been focused on limit cases, such as when the cell cycle duration is considered constant [6, 7, 9], or when DNA replication is omitted [10, 11]. Other studies have considered the effect of the cell cycle on protein fluctuations [5, 12–14]; the analysis in this case is simplified because unlike mRNA, protein lifetimes are very long and hence degradation is mostly due to dilution at cell division.

In addition, there are other factors beyond details of the cell cycle progression that can have a profound influence on transcript fluctuations. The symmetry of cellular division affects the number of transcripts in a cellular population. For instance, in a growing proliferating tissue, the continuous exponential appearance of young cells in a population introduces an asymmetry in the population cell age, favouring the proportion of cells at early stages of their cell cycle. This contrasts with the age structure of a homeostatic population where it is expected to find the cells equally distributed along their cell cycle [15, 16]. Since the average number of mRNAs in a cell increases with the time position in the cell cycle, we expect to observe larger mRNA content for the same type of cell in a homeostatic population compared to a growing population. Similar discrepancies arise when mRNA distributions measured from snapshots of a growing cell population are compared with the temporal tracking of the expression levels of a single cell over time, apparently contradicting ergodicity between single cells and the population. While this effect has been formalised mathematically [11], its relevance to the distributions of mRNA, or to the inference of different kinetic parameters, remains a conundrum.

In this paper we study the distribution of mRNA transcripts in single cells where expression can be bursty or non-bursty (both commonly observed, see for example [2]), with a cell cycle progression described as a number of stages having a stochastic duration. Our model also includes DNA replication and differentiates between population and lineage (single cell trajectory) measurements of the mRNA distribution. Keeping the framework relevant to the experimental inference of kinetic parameters, we aim to answer the following question: how important is the inclusion of cell cycle variability for predicting the statistics of stochastic mRNA expression? With this objective in mind, we derive and analyze expressions for the error made in different observables of transcript abundance when a deterministic cell cycle (one of fixed length) is considered instead of a stochastic one. Furthermore, we apply our results to a genome-wide expression dataset to address the magnitude of the error made in the inference of the transcription rate when mathematical models with different cell cycle details are employed.

## Model Description

We consider a general model of stochastic gene expression that takes into account cell cycle variability (for an illustration see Figs. 1a and b) with the following properties:

1. The cell cycle is divided into *N* stages. The duration of each stage *i* is exponentially distributed with a rate *k_i_*. This implies that the total cell cycle duration follows a hypoexponential distribution. Note that the number of stages, in general, will be larger than the number of cell cycle phases. Each biological cell phase can therefore be described as the composition of a given number of stages resulting in hypoexponentially distributed cell cycle phases consistent with recent experiments [17]. The number and duration of the different stages can be chosen by fitting the experimental cell cycle duration distribution.
2. The length of the mitotic phase is negligible and hence it is assumed to occur instantaneously after the end of the *N*-th stage. This leads to binomial partitioning of the mRNA between mother and daughter cells, where each individual molecule is allocated in either daughter cell with the same probability.
3. There is bursty or constitutive transcription of mRNA with rate *r_i_* of producing mRNAs per unit of time, and a decay rate *d_i_*. When the transcription is bursty, the burst size follows a geometric distribution with mean *β_i_*. All the parameters *r_i_*, *d_i_, β_i_* can vary depending on the stage *i* along the cell cycle.

**Figure 1:**
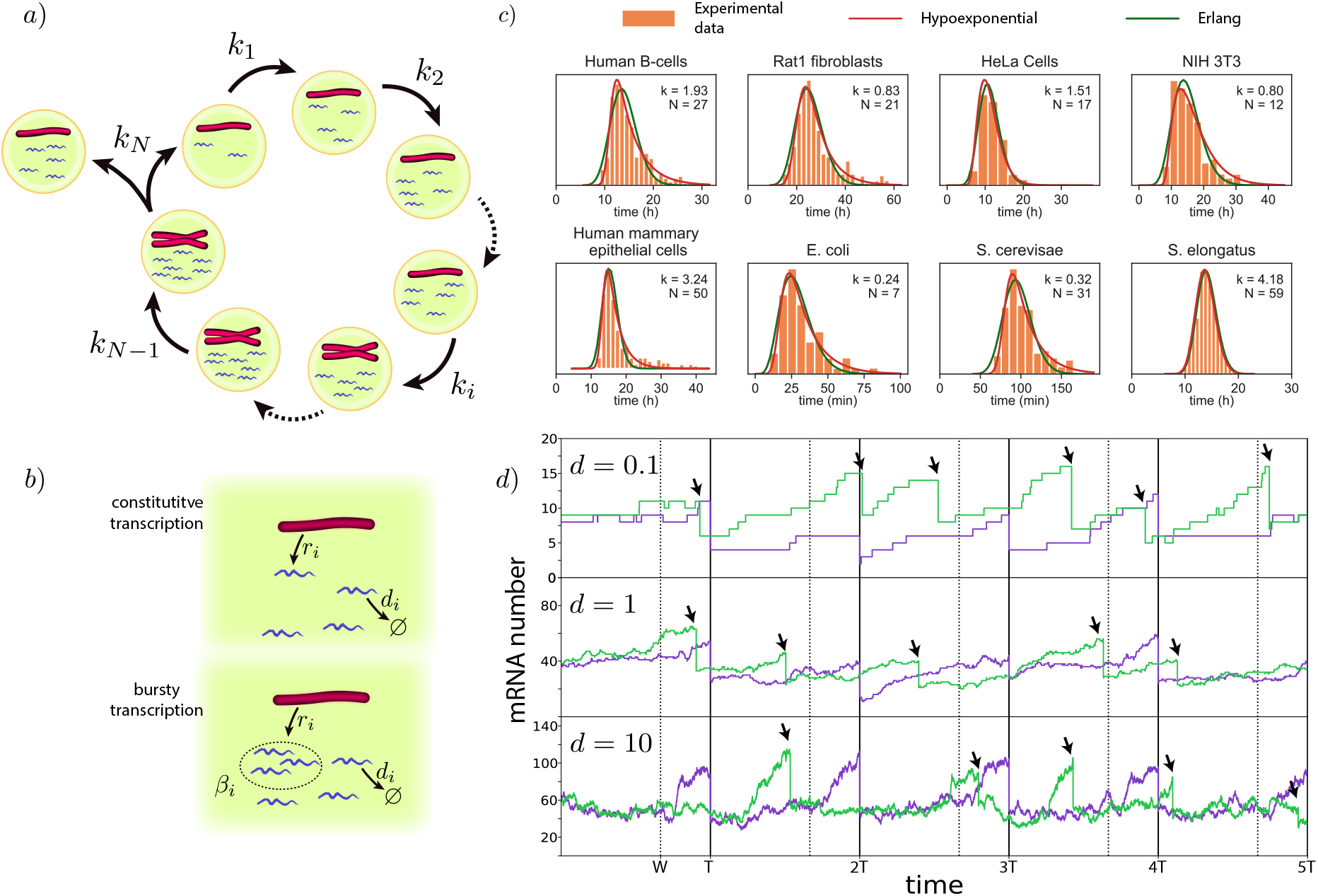
a,b) Schematic of the general model where mRNA dynamics take into account details of the cell cycle (a) including DNA replication of a gene, phase duration variability and bipartition of mRNA content at mitosis. During each cell cycle stage (b) mRNA dynamics is described as a production term (constitutive or bursty), and a linear degradation. c) The Erlang and hypoexponential distributions provide excellent fits to the experimental probability distributions of cell cycle durations of 8 different cell types. The parameters of the Erlang distribution (*k, N*) are shown on the figure. The sources of the experimental data, together with the parameters of the hypoexponential distribution (*k_i_, N*_1_, *k*_2_, *N*_2_) with *N*_1_ stages of rate *k*_1_ and *N*_2_ stages of rate *k*_2_ are: B-cells (11.7, 132, 0.26, 1) [21], Rat1 fibroblasts (0.15, 1, 2.4, 50) [22], HeLa cells (0.71, 4, 20.0, 110) [23], NIH 3T3 (0.24, 2, 11.4, 98) [19], mammary epithelial cells [24] (0.32,1,11.4,154), E. coli (0.88,16, 0.07,1) [25], S. cerevisae (0.04,1,1.4,115) [26], S. elongatus (4.18, 30,4.18,28) [27]. Fitted distributions correspond with the least-squares fit of the distance between the experimental histograms and the probability density functions using the Trust Region Reflective algorithm implemented in SciPy. d) Comparison of stochastic mRNA trajectories between a case where cell cycle duration is constant (purple) or stochastic (green), for different degradation rates *d*. Arrows indicate stochastic division times. Stochastic cell cycle simulations use the Erlang model with a production rate per chromosome equal to is *r* = 50*d* for a cell cycle with *N* = 4 stages, from which *W* = 3 stages occur prior to the gene replication (*w* = *W/N* = 3/4) indicated by dashed lines for the deterministic simulations.

This constitutes the general model studied in this manuscript. Detailing cell stage specific rates of transcription and degradation is particularly relevant since it will bestow our model with the ability to accurately describe the dynamic nature of mRNA transcription [3, 18]. In addition, for the sake of clarity of our analysis we will also consider a particular case of the general model:

1. All the cell stage rates are identical along the cell cycle *k_i_* = *k*. This implies that the total cell cycle duration follows an Erlang distribution. The number of cell cycle stages, in this case, can be easily determined from the best fit of an Erlang distribution to the experimental cell cycle duration [19]. In particular, the coefficient of variation (CV) of the Erlang distribution is equal to 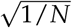.
2. The degradation rate of the mRNA is independent of the cell cycle stage *d_i_* = *d*.
3. There are *W* stages prior to DNA replication of the gene of interest and *N* – *W* stages postreplication. The production rate of mRNA is proportional to the DNA content of the cell at each stage without dosage compensation, being *r_i_* = *r* for *i* ≤ *W* and *r_i_* = 2*r* for *i* > *W*. We can consider replication to be instantaneous since the time of replication of the locus containing the gene of interest is much shorter than the total duration of the S-phase [20]. If transcription is considered to be bursty, the average burst size is constant along the cycle *β_i_* = *β*.

Since in this particular scenario the cell cycle duration follows an Erlang distribution, it will be referred hereon as the Erlang model to distinguish it from the general model. Despite the simplicity of the model, a fit of the Erlang distribution to 8 different types of eukaryotic and prokaryotic cells showed a very good fit, capturing the variability of cell cycle duration (Fig. 1c).

Stochastic simulations of the model can be used to study the effect of changing parameter values on the mRNA transcript number (Fig. 1d). Alternatively, we can study analytically the evolution of the probability *P_i_*(*n, t*) of finding a cell in stage *i* with *n* mRNAs at time *t* by using a Master Equation description, that for the general model with constitutive mRNA transcription (bursty case is detailed in the Appendix A) reads,

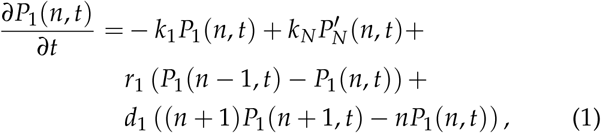

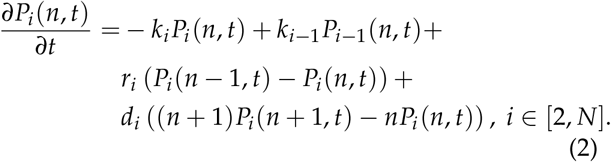

The first and second terms in these equations describe the exit from, and entrance to, the present cell cycle stage. The third term models transcription and the fourth term mRNA decay. Note that binomial partitioning during mitosis is explicitly taken into account by the second term of Eq. (1). This process implies:

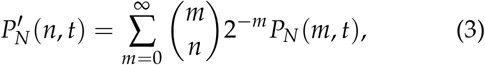

where we take the convention *m* choose *n* equals zero when *n* > *m*.

## Factorial Moments in Cyclo-stationary Conditions

Defining the generating function 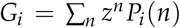, the Master Equations Eqs. (1)–(2) can be written as

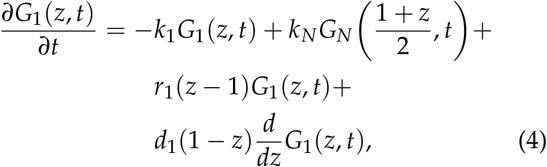

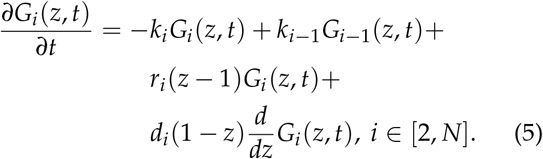

From the definition of the generating function it follows that the unnormalised *ℓ*-th factorial moment of the mRNA distribution in stage *j* is given by:

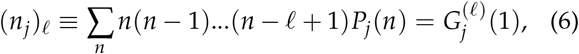

where the superscript (*ℓ*) means differentiating *ℓ* times. Enforcing cyclo-stationary conditions (steady-state for the mRNA distribution of each individual cell stage) by setting the time derivatives in Eqs. (4)–(5) to zero, differentiating *p* times the resulting equations and using the definition of the factorial moments above, we obtain:

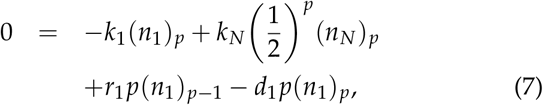

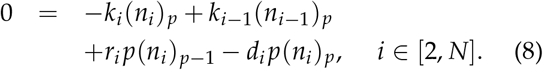

Eq. (8) can be brought into the form:

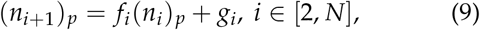

where we have used the definitions:

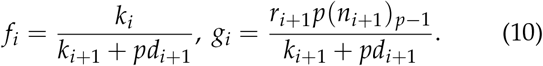

Since these are first-order non-homogeneous recurrence relations with variable coefficients, their solution can be written as:

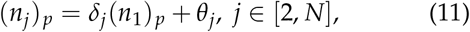

where we have used the definitions:

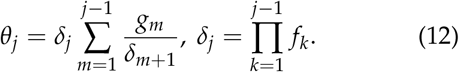

Solving Eq. (11) for (*n_N_*)_*p*_ and substituting in Eq. (7), after some simplification we obtain:

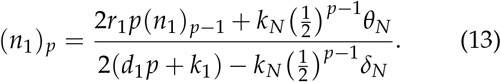

Note that the solution of the unnormalised *p*-th factorial moment depends on knowledge of the unnormalised *p* – 1-th factorial moment. Hence, because of this dependency, all factorial moments need knowledge of the zeroth order factorial moment (*n_j_*)_0_, which corresponds with the probability of finding the cell at stage *j*. By the definition of Eq. (6) we see that (*n_j_*)_0_ = *G_j_*(1). Setting *p* = 0 in Eqs. (7) and (8) one obtains:

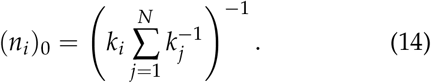

Hence summarising, Eqs. (11), (13), and (14) together give the solution to the unnormalised *p*-th factorial moment of the mRNA numbers in cell stage *j*. Note that to obtain the normalised *p*-th factorial moment one divides the unnormalised *p*-th factorial moment by 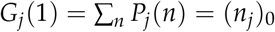.

The factorial moments for the general model with bursty transcription can be derived following the same steps. This procedure shows that the first factorial moment is equal to the constitutive case, whereas the factorial moments for higher orders in the bursty case are larger than in the constitutive case (see Appendix A).

## Lineage Measurements

The moments of the distribution can be used to compute the mRNA distribution statistics for different tissues. For instance, the mean number of mRNAs can be calculated as the average along the cell cycle stages of the expected number of mRNAs at each stage ((*n_i_*)_1_/(*n_i_*)_0_) weighted by the probability *π_i_* of finding a cell in a tissue at a certain stage *i*. Following this methodology, the expressions for the mean and the variance are,

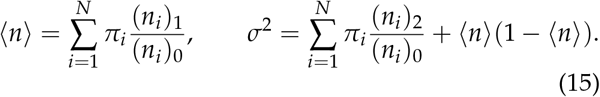

We will start our analysis studying the scenario in which the mRNA content of a single cell is tracked in time at regular intervals and, after division, the tracking keeps following only one of the daughter cells. This scenario is equivalent to the mRNA distribution of the cells forming a homeostatic tissue, where after each division one of the cells leaves the population, keeping constant the number of cells in the tissue [15, 16]. This scenario will be referenced in the text as the “lineage” case, to differentiate it from the mRNA distribution across a growing proliferating population of cells, which will be referred to as the “population” case. In the lineage case, the probability *π_i_* of finding a cell at the *i*-th cell cycle stage corresponds with (*n_i_*)_0_ (Eq. 14) being inversely proportional to the cell stage advance rate *k_i_*,

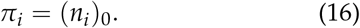

For the Erlang model this is *π_i_* = 1/*N*, and the explicit expression for the mean transcript can be obtained by introducing Eqs. (13), (14) and (16) in (15), obtaining,

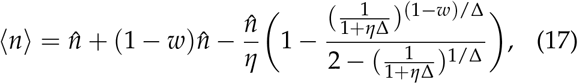

where for the sake of clarity we have written the expression in terms of the coefficient of variation of the cell cycle 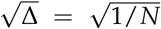. In addition, we have introduced the mean mRNA number in the absence of a cell cycle 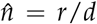, the fraction of the cell cycle before DNA replication of the gene of interest *w* = *W/N*, and the nondimensional parameter *η* = *dT* that compares the degradation timescale with the dilution timescale given by the average cycle duration *T* = *N*/*k* (see Table 1). Note that *η* is proportional to the ratio between the mRNA half-life *t*_1/2_ and the cell cycle duration *T* following *η* = *T*ln(2) / *t*_1/2_.

**Table 1:** Description of the different parameters used to describe the cell cycle, mRNA dynamics, and their relationship in the Erlang model. Parameters in shadowed rows can be derived from the rest of the parameters.

|  | Meaning | Erlang model |
| --- | --- | --- |
| $N$ | Number of cell stages | |
| $W$ | Cell stages prior to replication | |
| $k_i$ | Rate of advance of cell cycle stage $i$ | $k$ |
| $r_i$ | Transcription rate during stage $i$ | $r$ if $i \leq W$<br>$2r$ if $i > W$ |
| $d_i$ | mRNA degradation during stage $i$ | $d$ |
| $\beta_i$ | mean burst size during stage $i$ | $\beta$ |
| $w$ | Proportion of cell cycle before gene replication | $W/N$ |
| $\hat{n}$ | Stationary average mRNA number in absence of cell cycle | $r/d$ |
| $T$ | Average cell cycle duration | $N/k$ |
| $\Delta$ | Squared coefficient of variation of cell cycle duration | $1/N$ |
| $\eta$ | mRNA degradation relative to cell division rate | $dT$ |

The first term of Eq. (17) corresponds to the classical scenario without cell cycle. The second term of Eq. (17) introduces the effect of DNA replication for the case in which the mRNA degradation timescale is much shorter than the cell cycle length (*η* → ∞). Finally, the third term in Eq. (17) describes the contribution when mRNA degradation occurs at a comparable timescale to the cell cycle duration through the parameter *η* which has measured values in the approximate range 0.5 – 8.0 for a variety of cell types (see Table 2). This latter contribution increases monotonically with the cell cycle variability Δ (see Appendix B), and is minimal in the limit of Δ → 0 (deterministic cell cycle duration). In this deterministic limit Eq. (17) reduces to the simpler form

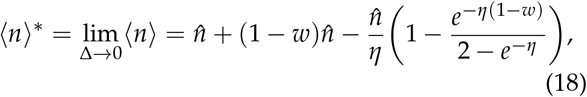

which agrees with a different calculation using deterministic rate equations (see Appendix C). Comparison of Eqs. (17) and (18) allows us to quantify the relative error *R* made in the expected number of mRNA when the cell cycle variability is not considered in the description of the mRNA dynamics,

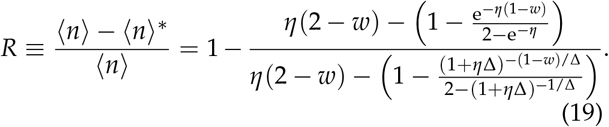

**Table 2:** Typical values of Δ, *w* and *η* for different cell types and their source. Values of Δ = 1/*N* are obtained from the Erlang fitting (see Fig. 1c). Range for *η* corresponds with the 99% CI of the *η* distribution for each genomic dataset.

|  | E. coli | S. cerevisiae | N1H3T3 |
| --- | --- | --- | --- |
| $\Delta$ | 0.14 [25] | 0.03 [26] | 0.08 [19] |
| $w$ | 0.15-0.50 [28] | 0.35-0.55 [29] | 0.40-0.80 [30] |
| $\eta$ | 0.41- 8.4 [31] | 0.89 - 15 [32] | 0.65 - 6.4 [33] |

Note that *R* is only a function of *η, w* and Δ, and therefore independent of the mRNA production rate (see Fig. 2a). The error is always positive (see Appendix B) and increases with the cell cycle time variability Δ, reaching its maximum for Δ = 1 (which is the maximum Δ attainable for an Erlang process since *N* ≥ 1). Similarly, since the expression for the first moment is identical in the bursty and constitutive cases (see Appendix A), the mean transcript number and its error are also independent of how bursty the transcription is.

**Figure 2:**
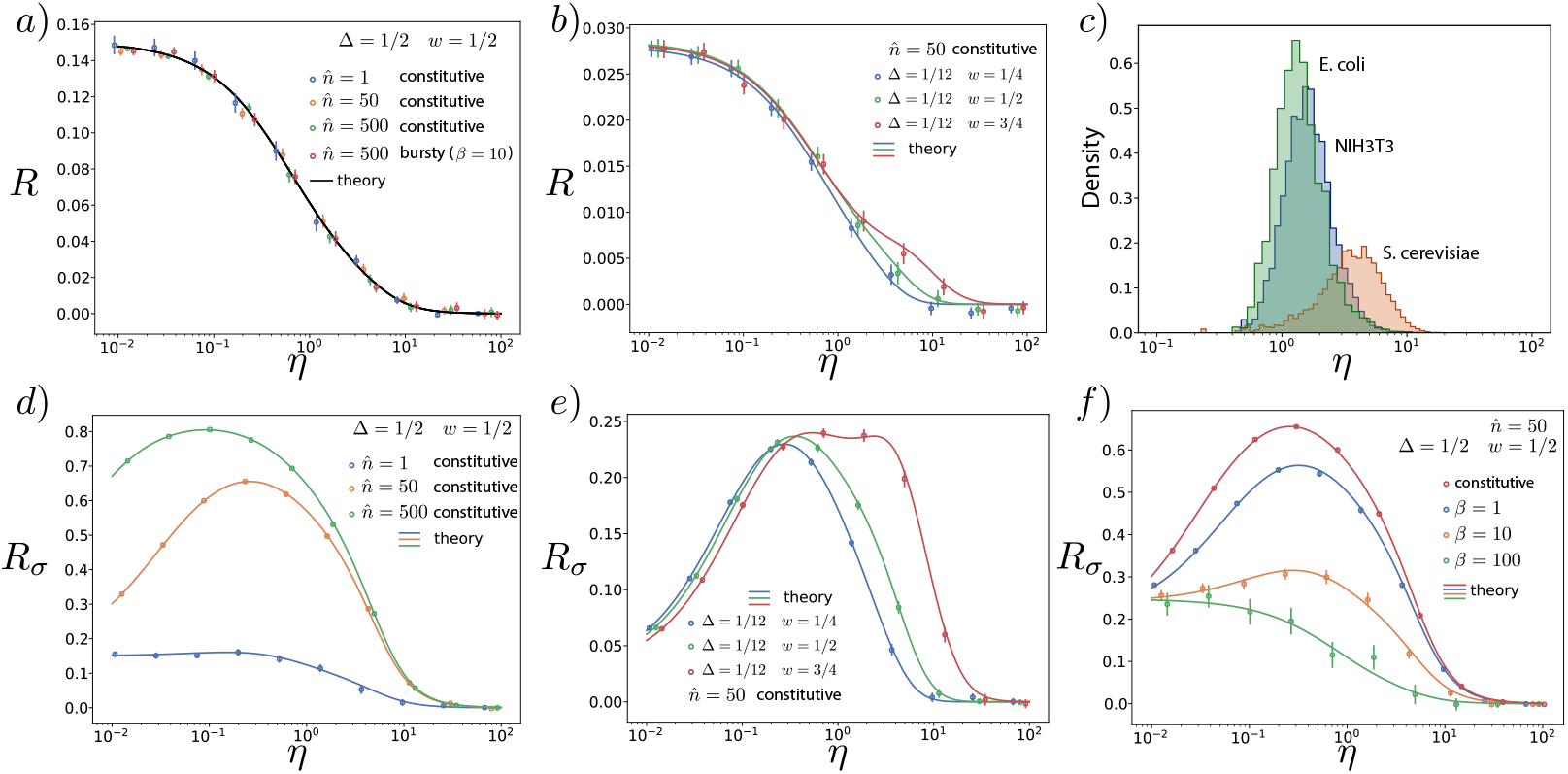
Relative error made in the average number of mRNAs (*R*) and its variance (*R_σ_*) when considering the cell cycle to be deterministic instead of Erlang distributed in a non-proliferating population or a lineage. Panels compare the theoretical results (lines) with stochastic simulations (circles). a,b) Relative error *R* of the mean number of mRNA. c) Genome-wide values of *η* for three different cell types. NIH3T3 mouse fibroblast data was obtained from [33]. Degradation rates for *S. cerevisiae* cultured in yeast extract peptone dextrose were obtained from [32] and its cell cycle duration from [34]. Stability data for the transcripts of *E. coli* cultured in Lysogeny broth were obtained from [31], while its cell cycle duration is taken from [35]. Averages are done over trajectories of duration *t* = 600*T* max(1/*dT*, 1 /*ρT*, 1). Panels a), b) and f) show averages over 50 trajectories for all conditions except for *n* = 1, which shows an average of 500 trajectories. Panels d) and e) show averages over 200 trajectories. Error bars indicate the standard error of the mean.

For a given cell type, the average time at which replication of a given gene occurs and the cell cycle duration variability can be considered constant (provided external conditions are not changed), and hence the value of the error *R* for different genes will be determined exclusively by *η*, which compares the mean cell cycle duration and mRNA lifetime, and can vary significantly from gene to gene [33]. The error decreases with *η* (see Figs. 2a,b), vanishing for *η* ≫ 1 corresponding with the scenario where mRNA lifetime is much shorter than the cell cycle duration. On the other hand, the relative error *R* is maximum for low values of *η*, describing the case of stable mRNAs for which degradation rates are much lower than the proliferation rate of the cell (analytical expressions for the mRNA distribution for this case can be found following the method described in [5]),

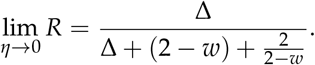

Interestingly, this maximal error depends on the properties of the cell cycle through *w* and Δ and it is maximised for intermediate levels of the DNA replication time 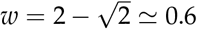, which is comparable to biological values of the relative duration of the G1 phase for N1H1 3T3 cells (see Table 2) [3, 30] (excluding cells which have arrested G_1_ phases), achieving a maximal relative error of *R* ≃ 15%, corresponding to Δ = 1/2 and 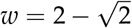.

The relative error can be more precisely estimated given data for specific types of cells. For example the cell cycle duration distribution in NIH 3T3 mouse embryonic fibroblasts has been described by an Erlang distribution with *CV*^2^ ≃ 1/12 (which implies *N* = 12 effective cell cycle stages) [19] and the G_1_ phase occupies roughly a fraction *w* = 0.4 of the cell cycle, indicating the position of the earlier transcribed genes during S-phase [30]. The maximum relative error *R* for these parameters lies around 3% (Fig. 2b), while for most of the transcriptome (*η* ∼ 1, see Fig. 2c) the relative error *R* ≃ 1%, indicating that in these cases the cell cycle duration variability can be ignored if the mean mRNA is all that we are interested in.

Making use of the second order moments of the distribution, we can extend the analysis to other statistic observables, allowing us to quantify the (relative) error in the variance, *R_σ_*, of mRNA fluctuations made when neglecting cell-cycle variability,

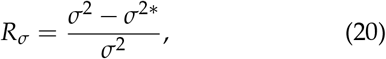

where *σ*^2*^ is the variance of the mRNA distribution in the deterministic cell cycle limit (Δ → 0). For the Erlang model, combining the expression for the variance (Eq. 15) with the factorial moments (Eqs. 13–14) we obtain an error for the variance *R_σ_* that is much larger than the one observed in the mean. Additionally, *R_σ_* does not have a monotonic dependence on the degradation rate, but is maximal for intermediate values of the degradation rate (*η* ∼ 1, see Figs. 2d,e,f). Interestingly, this region of values of *η* corresponds with most of the transcripts genome-wide for different species (see Fig. 2c). In particular, for the NIH 3T3 cells the error reaches *R_σ_* ∼ 25% (Fig. 2e), and can reach values as high as 80% for Δ = 1/2 (Fig. 2d). In contrast to the error in the mean, the error in the variance will depend on the transcription rate and the transcriptional burstiness. Analysis of *R_σ_* for the bursty model shows that *R_σ_* decreases with the burst size, reflecting that an increase in the variance due to the bursty gene expression reduces the relative impact of the contribution from cell cycle variability (Fig. 2f). Nevertheless, despite this reduction, the error *R_σ_* is still above 10% for many scenarios including both bursty and constitutive expression (Figs. 2d,e,f). Furthermore, in contrast to the error in the mean, *R_σ_* depends on the DNA replication position *w* in such a way that genes replicating later in the cell cycle (larger *w*) not only show larger errors but also for a broader range of degradation rates (see Fig. 2e).

## Population Measurements

When considering a proliferating population of cells, the continuous appearance of synchronised cells at an initial cell cycle stage establishes a different age distribution than the one derived in the lineage scenario (Fig. 3a). Specifically, after mitosis, one cell at stage *N* leaves the population to give rise to two cells at stage 1, enhancing the probability of finding cells in the population at initial stages of their cell cycle. The population values for the probability of observing a cell in the *i*-th cell stage *π_i_*, can be obtained by considering the evolution of the average number of cells in cell cycle stage *i* at time *t*, denoted by *C_i_*(*t*) [19, 36].

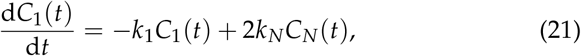

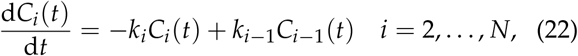

where the factor 2 in the first equation stands for cellular division: every time a cell divides (leaving stage *N*), two cells start at stage 1. In the lineage case this factor becomes 1. More generally, for cases with asymmetric division (after mitosis some cells leave the population with a certain probability) this factor 2 can be replaced by a factor *α* ∈ [0,2] [16]. While for Eqs. (21)–(22) the number of cells *C_i_*(*t*) will grow in time, the relative cell stage distribution in the population will eventually reach a steady-state for which we can write the ansatz *C_i_*(*t*)/*C*_1_(*t*) ≡ *λ_i_*. Specifically, for the Erlang case, introducing the definition of *λ_i_* in Eq. (22) yields the relationship

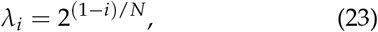

that gives the explicit values for the probability *π_i_* of observing a cell in the population at stage *i*,

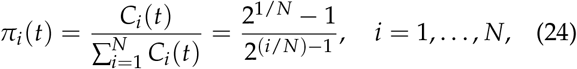

differing from the lineage stage distribution (Eq. (16), which for the Erlang case is constant (*π_i_* = 1/*N*). This discrepancy was confirmed by simulations (see inset of Fig. 3a).

**Figure 3:**
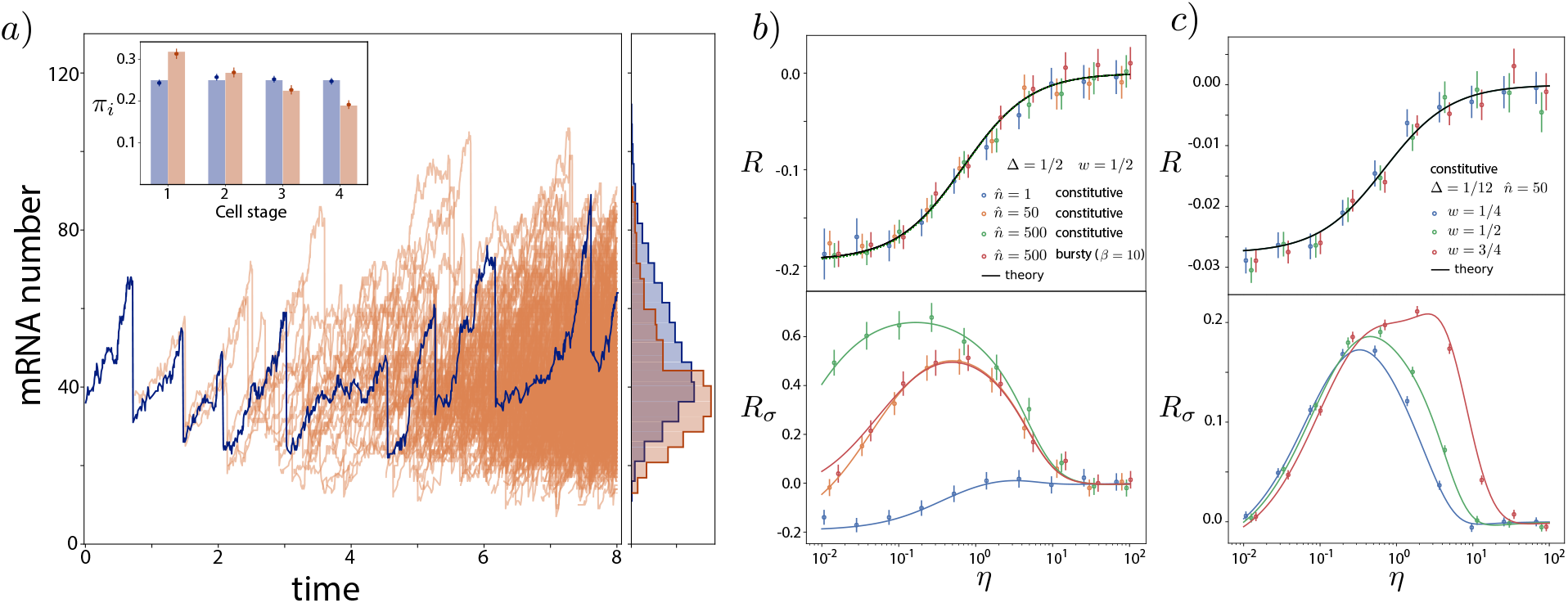
a) Comparison between the mRNA content over a single cell trajectory in time (blue) with the mRNA distribution of a proliferating population (orange). Histograms for both mRNA distributions (right) compare the average of 100 trajectory realizations with the snapshot of a single population at *t* = 7*T*. Parameters used are *T* = 1, *N* = 4, *W* = 2, 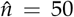, *η* = 1. Inset) Probability distribution *π_i_* of finding a cell at different cell cycle stages for single trajectory (blue) and a proliferating population (orange). Stochastic simulations for *π_i_* (circles) are compared with theoretical results (bars) obtained from Eq. (16) (that for the Erlang case is constant *π_i_* = 1/*N*) and from Eq. (24). b,c) Relative error made in the average number of mRNAs (*R*) and its variance (*R_σ_*) when considering the cell cycle to be deterministic instead of Erlang distributed in a growing proliferative population. Comparison includes theoretical results (lines) and stochastic simulations (circles). Simulations in b) show the average of 5000 snapshots at a time 10*T* and in c) the average of 25000 snapshots at a time 10*T*. Error bars indicate the standard error of the mean.

In addition to differences in *π_i_*, cells in the population case are also found more likely at earlier times inside each stage than in the lineage case. For the Erlang model, the distribution of times that each cell has been in its current cell stage follows an exponential distribution ∼ Exp(*k*2^1/*N*^) (see Appendix D and [5]). Given the Markovian nature of the process, this effect is equivalent to reducing *k* and having a faster cell advance through the cell cycle. Therefore, using the expressions for *π_i_* from Eq. (24), and the new effective rates of cell stage advance *k* → *k*2^1/*N*^ in Eqs. (11) and (13–14), allows us to obtain the factorial moments for population measurements. The mean number of mRNA in this scenario is

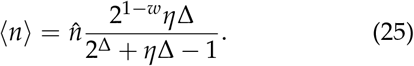

It is straightforward to show that 〈*n*〉 increases monotonically with Δ (similar to the lineage case). The exactness of Eq. (25) is confirmed by stochastic simulations in Figure 3. In the limit of a deterministic cell cycle, Eq. (25) reduces to the simpler form:

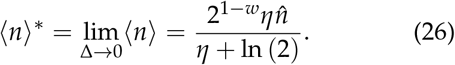

This agrees with a different calculation using deterministic rate equations (see Appendix C). Similar to the lineage case, this allows us to write explicitly an expression for the relative error *R* in the average number of mRNAs made when omitting the stochasticity of the cell cycle,

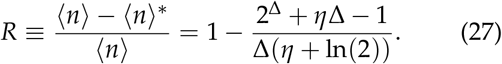

As in the lineage case, the error is a monotonic decreasing function of *η*, and increases with Δ reaching an error that is similar to the single cell case (*R* ≃ 20%) (see Fig. 3b and c). Nevertheless, in contrast to the lineage case, the error is negative, indicating that the expected number of mRNA decreases with the variability of the cell cycle duration. Strikingly, the error is independent of *w* and thus independent of the relative duration of G1 and G2 phases. Analysis of the error in the variance, *R_σ_*, results in similar observations to those of the lineage measurements, where *R_σ_* depends on the transcription rate and the transcription burstiness, resulting in errors much larger than *R* (*R_σ_* > 50%) that peaks at intermediate values of the degradation rate corresponding to the most frequent values of *η* measured genome-wide for different species (see Fig. 2c). As in the lineage case, the error *R_σ_* depends on the replication position during the cell cycle *w*, so genes replicating later in the cell cycle show larger errors for broader ranges of mRNA stability.

## mRNA Distribution Approximation

The exact mRNA distribution of our model is known only for some limit cases such as *η* → 0 [5]. Nevertheless, for more general realistic cases, we can use the moment derivation to reconstruct an approximate distribution. In particular, our analysis provides analytical expressions for the moments of the distribution at each cell stage *i*. Exclusively using the first moments, we can approximate the total mRNA population as a mixture of *N* Poisson distributions 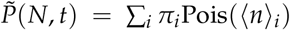, where the weights *π_i_* correspond to the probability of finding a cell at cell stage *i* obtained in Eqs. (16) and (24). Similarly, including the second moments, we can describe the probability as a mixture of negative binomial distributions 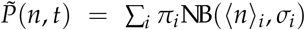, where each component 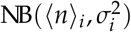 is a negative binomial distribution with mean 〈*n*〉_*i*_ and variance 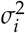 (see Eq. 15). Results for the lineage case, show that while the Poisson mixture failed to recover the distribution obtained from stochastic simulations in most scenarios (see Fig. 4a), the negative binomial mixture resulted in a very good prediction, able to recover the broad tails and bimodality of the mRNA distribution. In order to accurately assess the goodness of the reconstructed distribution, we computed the Kullback-Leibler divergence of the negative binomial mixture 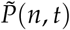 from the simulated exact distribution (see Fig. 4b). We observed that the approximation only fails for regimes with very unstable mRNAs that are highly expressed. On the other hand, the approximation improves for larger values of *N*, closer to experimental values for the cell cycle duration variability (*CV*^2^ = 1/*N* = 1/12) [19] (compare left and right panels of Fig. 4b). Comparison of the distributions for bursty expression and population measurements, using their corresponding moments and stages distributions, *π_i_*, yielded an even better approximation with values of the Kullback-Leibler divergence orders of magnitude lower than the lineage case (see Fig. 4c).

**Figure 4:**
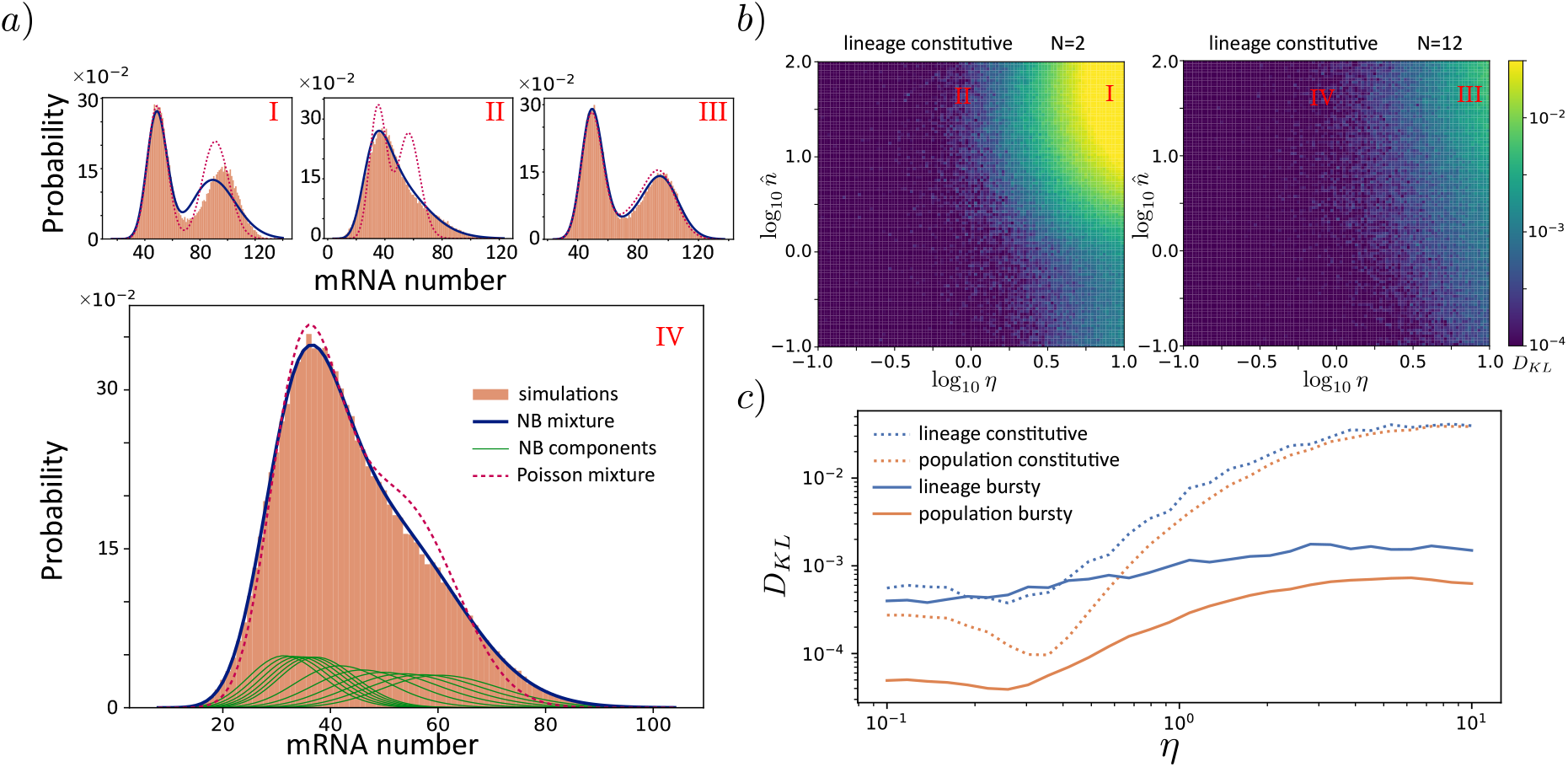
Approximation of the mRNA distribution as a mixture of negative binomials. a) Comparison between the approximations for the mRNA distribution (Poisson mixture and Negative Binomial mixture) with simulations for four different cases I (*N* = 2, *η* = 10, 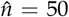), II (*N* = 2, *η* = 1, 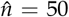), III (*N* = 12, *η* = 10, 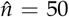), IV (*N* = 12, *η* = 1, 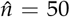). Case IV also includes the individual components of the NB mixture. b) Kullback-Leibler divergence of the NB binomial mixture from stochastic simulations for lineage distributions of constitutive mRNA expression. c) Comparison of the Kullback-Leibler divergence from simulations of the NB mixture for combinations of lineage/population measures and constitutive/bursty (*β* = 10) expression. For all the panels *w* = 1/2. Distributions for a) andb) result from trajectories over a time *t* = 6 · 10^4^*T* max(1/*dT*,1/*ρT*, 1), while for panel c) we used *t* = 6 · 10^6^*T/d* · max(1/*dT*,1/*ρT*, 1) for lineage measurements, and *t* = 20*T* for population measurements.

## Genome-wide Transcription Rate Inference Error

Our results so far have been focused on analyzing the errors that different models introduce on the mRNA statistics. Likewise, it is relevant to assess the error that different models introduce in the inference of biochemical parameters from experimental data. For this purpose we analyzed the genome-wide data from [33] and compared their transcription rate inference based on a lineage model with constant cell cycle and no replication (obtained by solving an expression equivalent to 〈*n*^*^〉 with *w* = 0 in Eq. (18), see Appendix F) against different models incorporating stochastic cell cycle duration (see Fig. 5 and Appendix F). Interestingly, since the average mRNA number is proportional to the transcription rate (see Eqs. 17 and 25), for given values of the cell cycle duration and gene replication, the relative error made when omitting cell cycle variation is a function depending only on the degradation rate through the parameter *η* (Fig. 5a). Since [33] reported no correlation between mRNA stability and transcription rate, this resulted in an absence of correlation between the error and the speed at which genes are transcribed (Fig. 5a). Additionally, in agreement with the error of the average mRNA number *R*, the error in the transcriptional rate estimate increases with the stability of the mRNA. When comparing the error expected for different models, small errors were observed for the lineage case with no DNA replication (See Fig. 5b). Nevertheless, for more realistic scenarios where the error is evaluated for a growing population case with DNA replication [33], more than 90% of the genes detected underestimate the transcription rate with an error bigger than 10% (see Fig. 5b).

**Figure 5:**
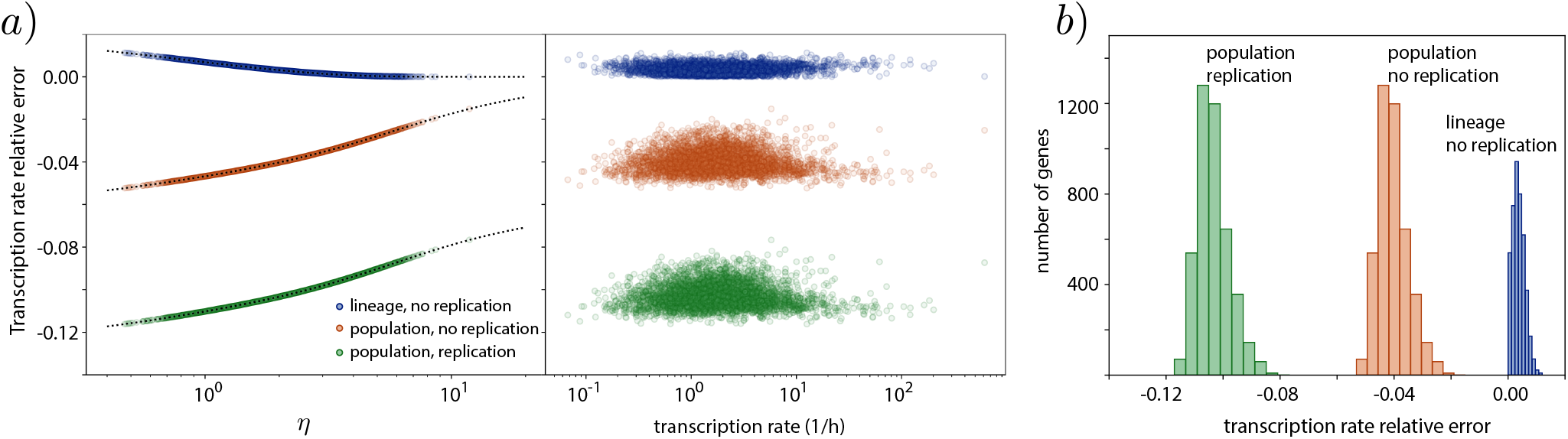
Relative error of inferred mean transcription rates with deterministic models at a genomic scale. a) Relative error in the inferred transcription rate for the 5028 genes reported in [33], as a function of the relative degradation rate *η* and the reported transcription rate. The relative errors are calculated between different models with stochastic cell cycle and the deterministic cell cycle model without replication used in [33] (see Appendix F). Discrepancies between models are a function of *η* (dotted line). b) Histogram showing the number of genes for different levels of transcription rate error for three different stochastic cell cycle models. The stochastic cell cycle used is an Erlang model with *k* = 0.145*h*^−1^ and *N* = 16 using the reported cell cycle duration and variability in [33] (see Appendix F), while replication is considered to occur at the middle of the cell cycle (*w* = 1/2).

## Discussion

Most of the models employed to study gene regulation ignore the effect that a detailed stochastic cell cycle description has on gene expression. The model and methodology developed in this paper not only allows one to analytically evaluate the role of features such as cell duration stochasticity or DNA replication in the transcript population but also provides a straightforward way of discriminating the scenarios for which such details are relevant for the description of the system. This is of paramount importance when mathematical models are used to infer parameters from experimental data, where the precision of the information demands the use of the right level of abstraction [37].

Specifically, this approach contrasts with alternative strategies that either ignore cell cycle effects or fit mRNA populations to arbitrary population mixtures, impeding the inference of mechanistic information of the transcriptional parameters. This is of particular relevance for current data analysis where mRNA labelling techniques give access to mRNA abundance distributions in populations of cells. In order to extract mechanistic information of the transcriptional process from these distributions, it is paramount to link the details of the distribution to the properties of the different biomolecular mechanisms [3, 38]. While in this paper we analyzed the error in the transcription rate estimation due to neglecting cell cycle variability and replication, future work will address how taking into account such details may also affect the inference of other biochemical parameters such as gene activation and deactivation rates, or the mean burst size. The necessity of such a study becomes apparent from the mRNA distributions obtained, which can be approximated accurately by negative binomials in scenarios with constitutive gene expression, challenging the common practice to use negative binomial distributions as a signature of bursty transcription [1, 38, 39]. Experimental validation of these predictions will require inference of parameters using data obtained by live imaging techniques, such as labelling with mRNA aptamers such as MS2, Mango, or Peppers [40–42] capable of measuring individual mRNA dynamics. Specifically, trajectory data provided by live imaging techniques can be used together with the moments provided by our stochastic cell cycle length theory to infer the parameters of the model using Bayesian methods (such as ABC) [37]. Contrasting the resulting parameter posterior from a stochastic cell cycle model with those obtained from performing the same analysis under the assumption of a fixed cell cycle length can highlight the relevance of cell cycle variability for the estimation of transcription and other biochemical parameters.

One of the major assumptions used in the bulk of the paper is that all the stages of the cell cycle have the same rates of mRNA production per allele, burst size, and degradation. Nevertheless, our analysis allows the incorporation of different rates of production and degradation in the expression for the moments of the general model (Eqs. (11–14). Such an accurate description will be of paramount importance to understand the effect of stochastic cell cycle duration in genes related to cell-cycle progression [43–45] or dosage compensation along the cell cycle [3, 46]. Furthermore, additional experimental stochastic details of the different cell cycle phases can be incorporated by choosing accordingly the rates and number of stages of the general model [47].

Future extensions of the model can focus on incorporating more detailed descriptions of the stochastic dynamics of the different processes. These could include more realistic assumptions about how molecules are partitioned at cell division to effectively account for specific segregation mechanisms [8]. Another possible extension could describe transcriptional bursts by considering promoter switching dynamics explicitly [48] allowing us to investigate the effect of multiple promoter states on the mRNA dynamics [49].

Finally, further extension of the model should include protein regulation of mRNA abundance. Particular mechanisms include nuclease dynamics controlling mRNA turnover along the cell cycle [43, 50], or cyclin-dependent kinases controlling cell-cycle advance [51]. In addition, incorporating the methodology developed in this paper to gene regulatory networks will provide a route to better understanding the stochastic details of the expression of genes with dynamics that can change in a timescale comparable to the cell cycle duration, such as circadian clock related genes [52, 53]. This is of special relevance in embryonic development, where the details of intrinsic noise are known to play a major role in the formation of spatial domains of gene expression in the patterning of embryonic tissues [54–56].

## Acknowledgments

We thank James Briscoe for feedback on the manuscript. R.P.-C. acknowledges support from the UCL Mathematics Clifford Fellowship. R.G. acknowledges support from the Leverhulme Trust (Grant No.RPG-2018-423). C.B. acknowledges the Clarendon Fund and New Col-lege, Oxford, for funding.

## Appendix

## A Bursty mRNA transcription model

Considering the general model, we can introduce bursty mRNA transcription as a reaction with a burst rate *v_i_* at cell stage *i*. The number of mRNAs *ℓ* produced in a burst at cell stage *i* follows a geometric distribution *ξ_i_*(*ℓ*) with average number *β_i_* of transcripts produced per burst i.e. an average rate *r_i_* = *v_i_β_i_* of mRNAs produced per unit of time [57]. The explicit geometric probability distribution follows

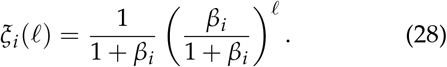

The resulting Master Equation reads,

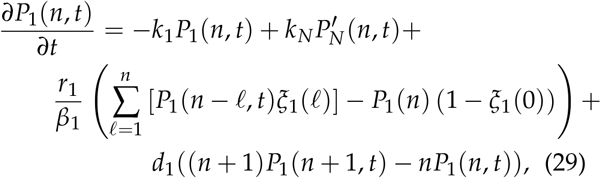

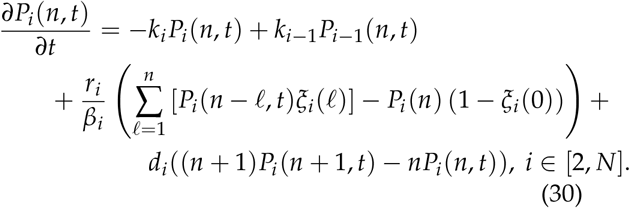

As in the constitutive case, we can use these system of differential equations to obtain the steady state factorial moments of the distribution by introducing the generating function 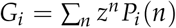. In particular the terms corresponding to bursty transcription follow the sum,

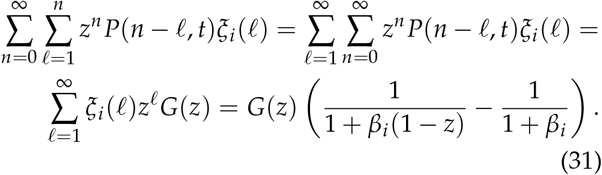

This results in the system of differential equations,

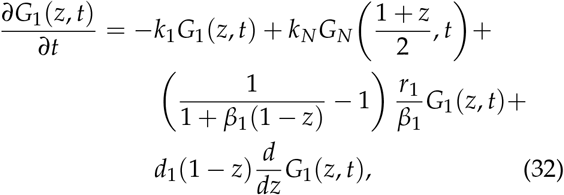

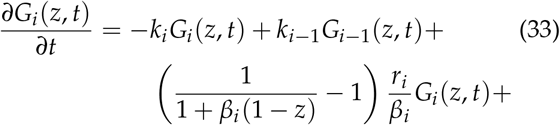

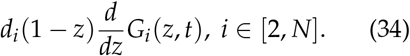

Enforcing the steady-state by setting the time derivatives in Eqs. (32) and (33) to zero, differentiating *p* times the resulting equations and using the definition of the factorial moments (*n_i_*)*k* we obtain:

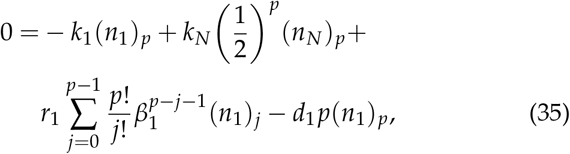

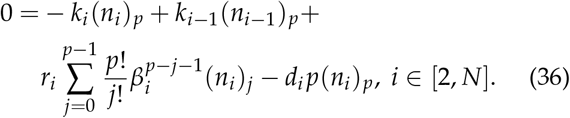

The normalization of the factorial moments (*n_j_*)_0_, obtained for *p* = 0 with the normalization condition 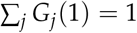, is the same as in the constitutive case, and only depends on the cell stage advance rates

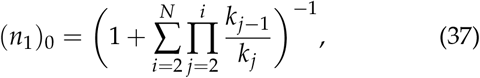

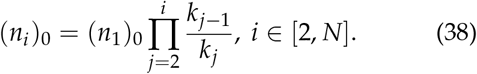

Similarly to the constitutive case, equation (36) can be written in the form,

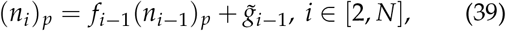

where we have used same definition for *f_i_* as in the constitutive production case, but 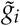 replaces *g_i_*, which instead of depending on the immediately lower order factorial moment *p* – 1, depends on all the moments lower than *p*:

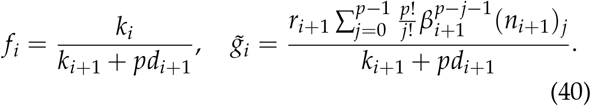

These first-order non-homogeneous recurrence relations have the solution

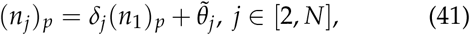

where we have used the definitions:

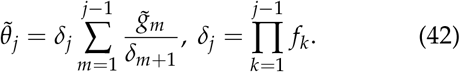

Substituting this solution in Eq. (35) we obtain

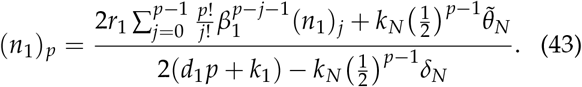

Comparing these results we can immediately see that the expected value of mRNAs is the same in the bursty case and the constitutive case considering the same average rate of mRNA production at cell stage *i*: *r_i_* = *v_i_β_i_*. Differences arise for higher moments. In particular all the factorial moments of the bursty scenario with *p* > 1 are larger than the factorial moments of the constitutive case since,

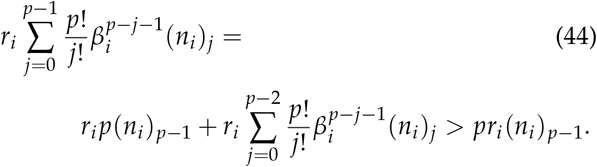

## B Monotonic dependence of mean mRNA on the coefficient of variation of the cell cycle duration for lineage observations

By Eq. (17), we have for *η* > 0

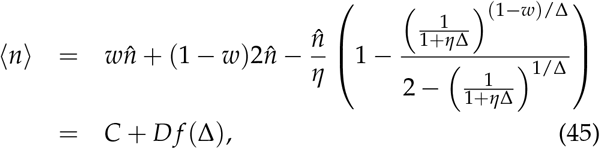

where *w* is a fraction, *C, D* are constants (*D* is positive) and

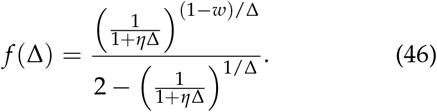

If we define *x* = (1 + *η*Δ)^1/Δ^ we note that since Δ > 0 we have *x* ∈ (1, *e^η^*) and also *x*(Δ) is monotonically decreasing, which follows from

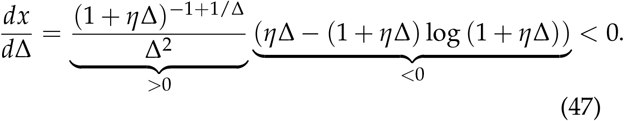

We then note that using this transformation we get:

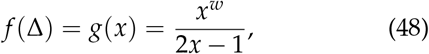

which satisfies

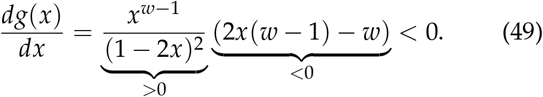

Using this we find

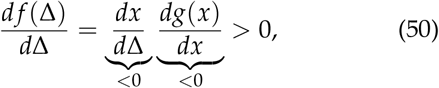

which proves strict monotonicity of 〈*n*〉 as a function of Δ > 0.

## C Alternative derivation of Eqs. (18) and (26) from deterministic rate equations

Consider a cell cycle of fixed duration *T* with replication (and consequent doubling of transcription) occurring at time *τ* = *wT* (where *w* is a fraction). If the transcription rate before replication is *r*, the mRNA decay rate is *d* and *n*(*t*) is the deterministic estimate for the mean number of mRNA molecules at time *t* then a deterministic model for this process is:

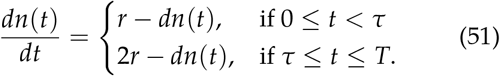

In the cyclo-stationary limit, binomial partitioning (when cell division occurs at the end of the cell cycle) leads to the boundary condition 2*n*(0) = *n*(*T*). Note that *t* in this context means the cell age and not absolute time and hence it can only vary between 0 and *T*. Solving these differential equations we obtain the solution:

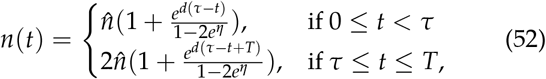

where *n* = *r/d*. Let *f*(*t*)*dt* be the probability of observing a cell of age between *t* and *t* + d*t* where d*t* is an infinitesimal time interval. It then follows that:

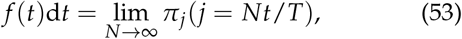

where *π_j_* is the probability of observing a cell in cell cycle stage *j*. Note that since Δ = 1/*N*, the limit of *N* → ∞ at constant *T* is the same as the limit of Δ → 0. Since *T* = *N*/*k*, in this limit we have infinite cell stages *N* advancing with an infinite rate *k* i.e. the cell spends an infinitesimal small time d*t* = 1/*k* at each stage. Knowing that *π_i_* = 1/*N* for lineage measurements we have

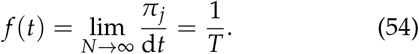

For the population case we substitute *i*/*N* = *t*/*T* in Eq. (24) take limit of large *N* and finally use *N* = *kT* to obtain,

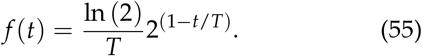

Note that both Eqs. (54) and (55) are well known and have been in common use for more than 40 years [58]. Finally we obtain the mean number of mRNA averaged over the cell cycle 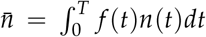. For the lineage measurements this yields

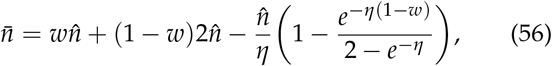

whilst for the population measurements we obtain

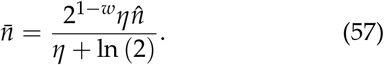

These expressions agree exactly with Eqs. (18) and (26) which were derived from a Master Equation approach in the limit of zero variability in the cell cycle duration for the case of lineage and population measurements, respectively.

## D Derivation of the distribution of cell stage durations in population measurements

We let *C_i_*(*t*, τ) denote the number of cells in a population that are in cell stage *i* at time *t* that have been in that cell state for a duration *τ*. After a small time duration *δ* all the cells will either advance to an age *τ* + *δ* or advance to the next cell stage. Therefore we can write the conservation equation

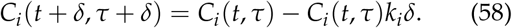

Assuming that there is a stationary distribution for the stage age of the cell population at a stage *i, p_i_* (*τ*), we can write *C_i_*(*t,τ*) as *C_i_*(*t,τ*) = *C_i_*(*t*)*p_i_*(*τ*). Where *C_i_*(*t*) is the number of cells at cell stage *i*. Introducing this factorization of *C_i_*(*t, τ*) in Eq. (58), and taking the limit *δ* → 0, we get the relationship,

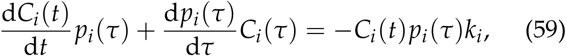

where we have used the chain rule to compute the derivative of d*N_i_*(*x,x*)/d*x*|_*t,τ*_. Since the probability of finding a cell in a certain stage *i, π_i_*, is constant in time, the number of cells at a given stage has to grow with the same rate as the population, therefore d*C_i_*(*t*)/d*t* = *KC_i_*(*t*), where *K* is the growth rate of the population. Introducing this equality in Eq. 59, we get an equation for *p_i_*(*τ*).

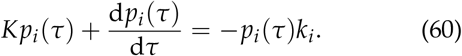

That gives,

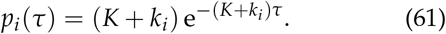

In the Erlang distributed model, *k_i_* = *k* is constant, and the rate of growth of the population can be calculated from the conservation equation for the total number of cells *C*(*t*),

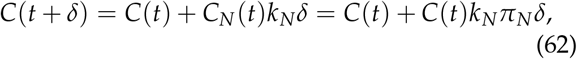

where *N* is the number of stages of the cell cycle. From this equation, we obtain that the rate of exponential growth of the population is *K* = *k_N_π_N_*. Using the value of *π_N_* from Eq. (24), we obtain that for the Erlang model, the stage age distribution of cell cycle stage *i* is

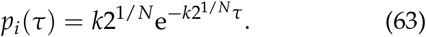

## E Computational analysis

The simulations for the general cell cycle model (including Erlang distributed times), were made using a custom made publicly available Gillespie algorithm where cell cycle stages are treated as one extra reaction (https://github.com/2piruben/langil/tree/master/examples/CellCycleVariability). After the last stage of the cell cycle is completed, the cell cycle time is reset and the number of mRNAs is reduced by sampling a binomial distribution *B*(*n*, 1/2) where *n* is the number of mRNAs before cell division (see Algorithm 1).

### Algorithm 1 General model with stochastic cell cycle with constitutive expression

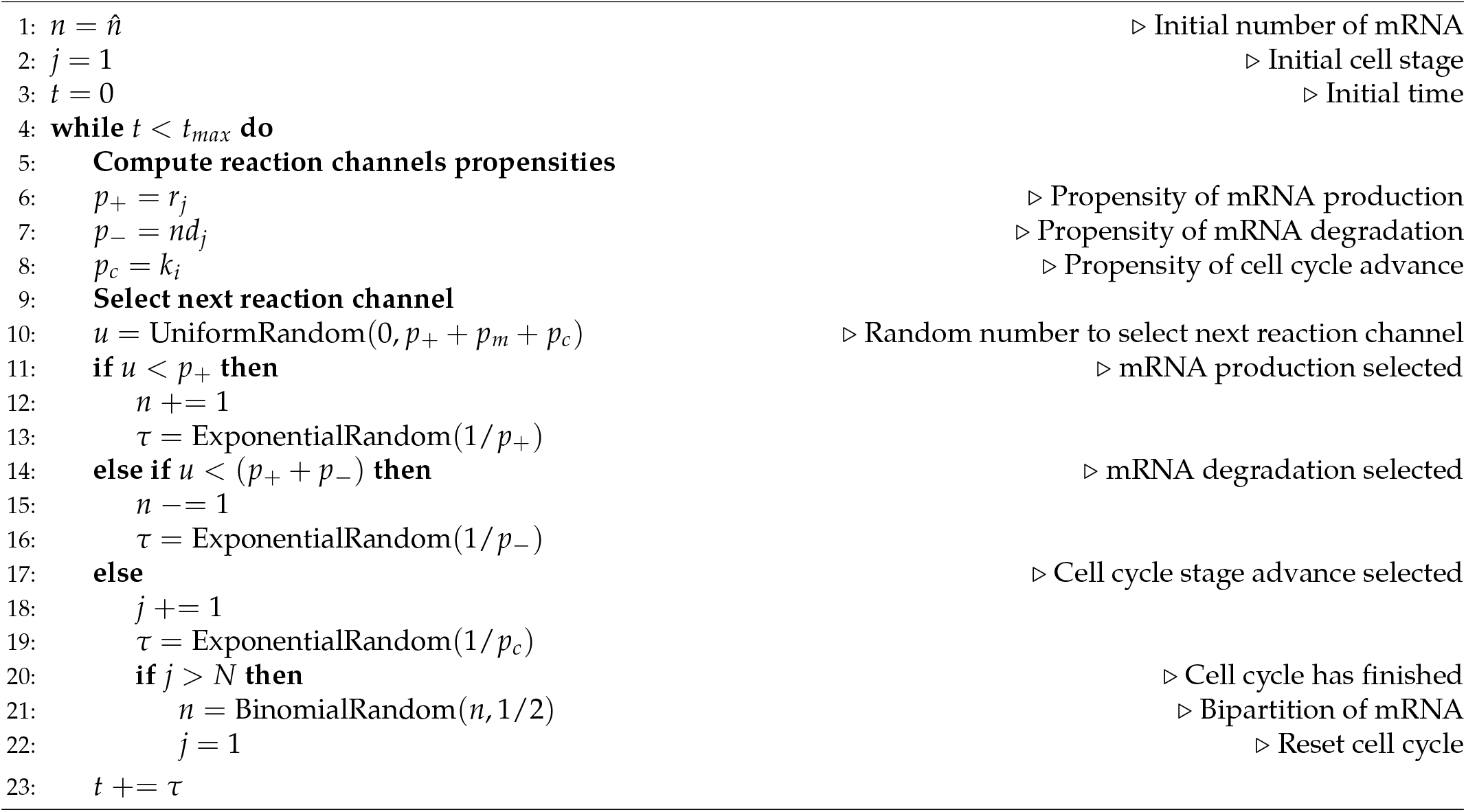

On the other hand, to simulate a cell cycle where the different stages have deterministic duration, the Gillespie algorithm has been modified to take into account if a deterministic cell stage change would take place before the next stochastic reaction time (see Algorithm 2). The rest of the details of the algorithm are the same as in the general cell cycle model.

### Algorithm 2 Deterministic cell cycle model

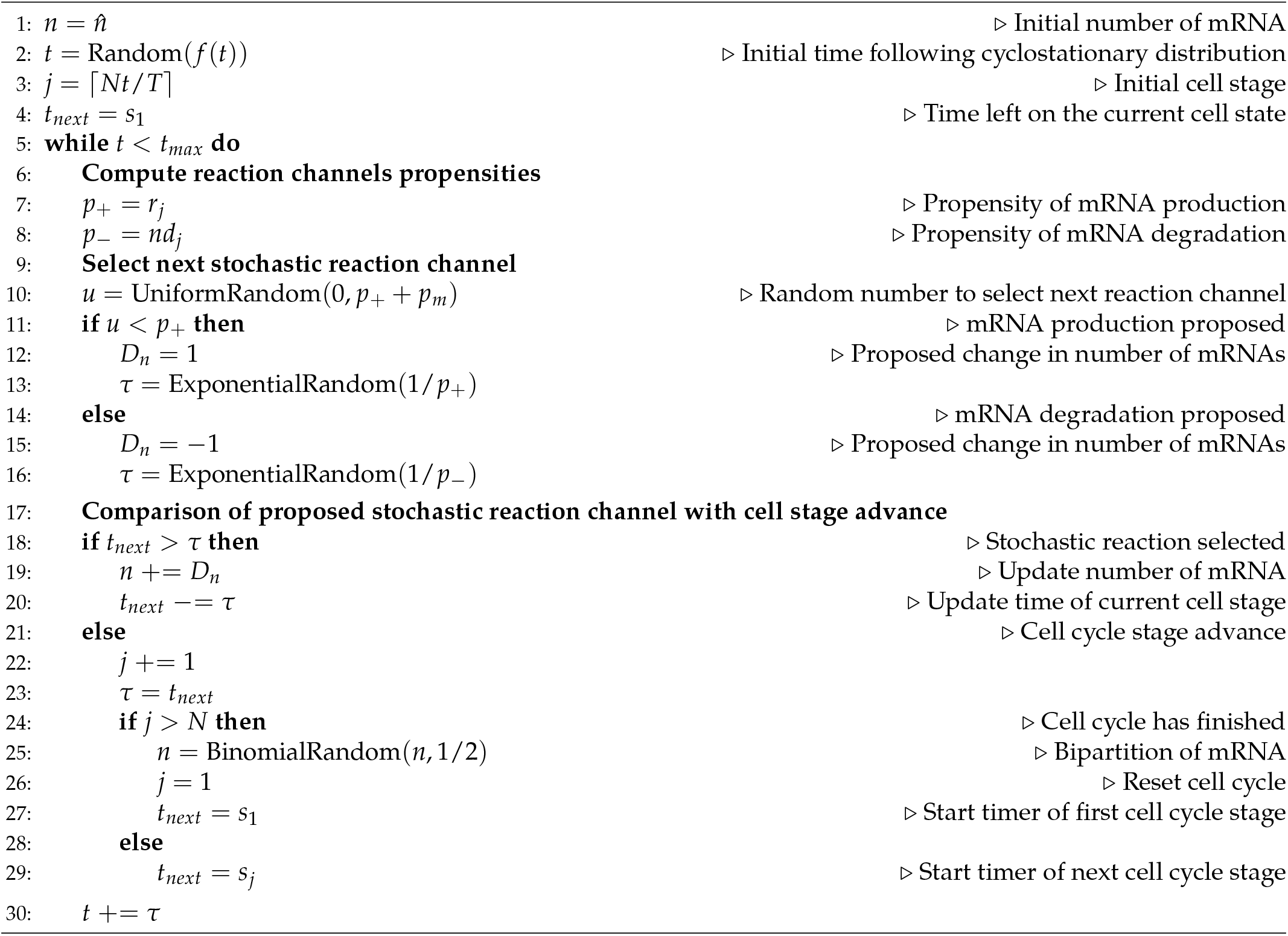

To obtain statistics from lineage measurements, each trajectory was sampled by choosing evenly distributed time points. For population measurements, several simulations are run in parallel, one for each cell. After each cell division event, a new cell is introduced in the simulation containing the remaining mRNA from the binomial partition of the mother cell. In order to achieve a steady state behaviour with deterministic cell cycles it was necessary to initiate each replicate following the corresponding age distribution (Eq. (54) or Eq. (55)). Statistics from the population measurements are calculated across all the cells at a particular time snapshot.

## F Inference of transcription rates and error calculation

Using the expression for the average number of mRNAs in the lineage measurements given by Eq. (17), we can write the transcription rate parameter *r* for the Erlang model as a function of the average number of mRNAs observed and the rest of the parameters of the model,

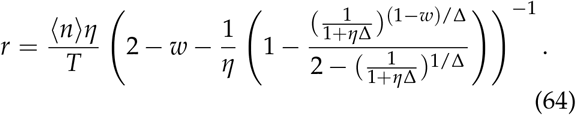

Similarly, we can write an expression for *r* for the population case using Eq. (25),

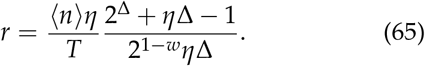

We can use both Eqs. (64) and (65) to obtain the average transcription rate *r* along the cell cycle for lineage or population cases,

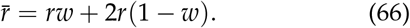

In the limit of a deterministic cell cycle duration and no replication 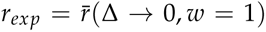 we recover the expression used in [33], which returns a value of transcription rate for each gene given the measured decay rate and average number of mRNA transcripts. By contrast, in order to compute 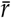 in a general case we need to evaluate the cell cycle duration variability Δ. The cell cycle duration reported in [33] was 19.9h < 27.5h < 33.6h, and hence we choose a standard deviation of (33.6-19.9)/2 = 6.85h. The corresponding number of effective states *N* can be obtained from the coefficient of variation of the cell cycle length *N* = 1/Δ = 1/*CV*”^2^ ≃ 16.

Introducing the calculated value of Δ in Eqs. (64–66), we can evaluate 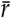 for lineage and population cases for different DNA replication positions along the cell cycle. In the text we study a case without replication *w* = 1 and a case with replication at the middle of the cell cycle *w* = 1/2.

In order to evaluate how our predictions differ from the reported transcription rates *r_exp_*, we compute the relative error *ε*

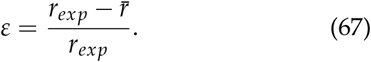

Note that since 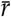 is linear in the average transcript number 〈*n*〉, the resulting error is independent of 〈*n*〉. Therefore, differences in the error *ε* among the different genes reported in [33] will only depend on their degradation rate.

